# Mitochondrial Complex I Modulator Restores Network Resilience in Advanced Alzheimer’s Disease Through Metabolic Reprogramming

**DOI:** 10.64898/2026.06.14.732179

**Authors:** Esraa Gabal, Thi Kim Oanh Nguyen, Tetiana Kovalenko, Huanyao Gao, Noa Rappaport, Cory Funk, Priyanka Baloni, Eugenia Trushina

**Affiliations:** School of Health Sciences, Purdue University, West Lafayette, IN 47907, USA; Purdue Institute for Integrative Neuroscience, Purdue University, West Lafayette, IN 47907, USA; Department of Neurology, Mayo Clinic, Rochester, MN 55905, USA; Department of Molecular Pharmacology and Experimental Therapeutics, Mayo Clinic, Rochester, MN 55905, USA; Institute for Systems Biology, Seattle, WA 98109, USA

**Keywords:** Alzheimer’s disease, mitochondrial complex I modulators, APP/PS1 mice, lipid metabolism, energy homeostasis, mitochondrial dysfunction, neuroprotection, genome-scale metabolic models

## Abstract

Mitochondrial dysfunction and lipid dysregulation are among the earliest abnormalities in Alzheimer’s disease (AD), yet their mechanistic interplay and therapeutic potential remain poorly understood. Here, we investigated whether restoration of mitochondrial function can reverse metabolic dysfunction and promote resilience in advanced-stage AD. Female APP/PS1 mice were treated with the brain-penetrant mitochondrial complex I (mtCI) modulator CP2 beginning at 19 months of age, when pathology and cognitive deficits were well established. To define the metabolic mechanisms underlying therapeutic response, we developed *iMiceBrain*, the first brain-specific genome-scale metabolic model of the mouse brain, and integrated transcriptomics, targeted metabolomics, lipidomics, and metabolic network analyses. CP2 treatment broadly reprogrammed AD-associated molecular signatures and restored pathways involved in mitochondrial function, glucose utilization, lipid metabolism, synaptic activity, and cellular stress responses. Metabolic modeling identified enhanced mitochondrial substrate flexibility, activation of fatty acid utilization, restoration of pyruvate dehydrogenase flux, and normalization of cholesterol metabolism as key features of the therapeutic response. Lipidomic analyses further demonstrated correction of disease-associated alterations in cholesteryl esters, phospholipids, and sphingolipids. Together, these findings demonstrate that mild mtCI modulation restores metabolic resilience by coordinating mitochondrial and lipid metabolism, establishing it as a disease-modifying therapeutic strategy for AD.

## Introduction

Alzheimer’s disease (AD), the most prevalent form of dementia, is characterized by progressive cognitive decline driven by a multifactorial etiology comprising genetic susceptibility, aging, neuroinflammation, and metabolic dysfunction. The extracellular accumulation of amyloid-β (Aβ) and intracellular aggregation of hyperphosphorylated tau (pTau) are widely recognized as canonical pathological hallmarks of AD. These features are associated with synaptic failure and neuronal loss, while age and the apolipoprotein E4 (APOE4) genotype represent the strongest risk factors for disease susceptibility. Multi-omics studies of post-mortem human brain tissue and biofluids reveal that AD progression involves disease-stage- and sex-specific alterations across interconnected metabolic and signaling pathways^1,2^. Among these alterations, lipid dysregulation and mitochondrial dysfunction are among the earliest detectable abnormalities, preceding overt Aβ and pTau pathology^2–5^. Emerging evidence indicates that mitochondrial dysfunction and lipid dysregulation form a self-reinforcing pathological network in AD, whereby impaired bioenergetics disrupt lipid homeostasis, while aberrant lipid metabolism further compromises mitochondrial function and cellular resilience. Lipid abnormalities in AD include disrupted cholesterol homeostasis, altered phospholipid and sphingolipid composition, impaired lipid droplet dynamics, reduced plasmalogens, and accumulation of lipid peroxidation products, collectively compromising membrane integrity, synaptic function, mitochondrial activity, and myelin stability^4,6^. Consequently, lipid disturbances are increasingly viewed not merely as secondary consequences of neurodegeneration but as active contributors to disease pathogenesis^7,8^. Maintenance of lipid homeostasis requires coordinated interactions between mitochondrial metabolism and cellular lipid-handling pathways. Mitochondria provide the energetic and biosynthetic support necessary for cholesterol trafficking, fatty acid oxidation, lipid droplet turnover, and membrane remodeling. Failure of these processes contributes to neuroinflammation, impaired neuron-glia communication, and progressive synaptic dysfunction in AD.

Beyond ATP production, mitochondria function as central signaling hubs that coordinate calcium homeostasis, lipid metabolism, redox balance, inflammation, epigenetic programs, and adaptive stress responses^9^. Perturbations in tricarboxylic acid (TCA) cycle intermediates, including acetyl-CoA, citrate, α-ketoglutarate, succinate, and fumarate, influence histone methylation and acetylation, linking mitochondrial metabolism to epigenetic regulation of gene expression, consistent with the global DNA hypomethylation observed in AD brains^10^. Mitochondria also interact extensively with the endoplasmic reticulum to coordinate lipid trafficking, calcium signaling, autophagy, immune responses, and cell survival^11^, processes that become progressively disrupted during AD progression.

The causal contribution of mitochondrial dysfunction to AD pathogenesis is increasingly supported by both experimental and human evidence. We recently demonstrated that a ∼50% reduction in mitochondrial complex I (mtCI) activity, achieved by deleting the *Ndufs4* subunit, induces sex-specific AD-like transcriptomic signatures in the mouse brain that closely mirror those observed in familial AD mouse models and patients with late-onset AD (LOAD)^3^.

Dysregulated pathways included lipid metabolism, inflammatory signaling, synaptic function, and epigenetic regulation, with particularly pronounced effects in females. These findings are consistent with the mitochondrial cascade hypothesis, which proposes that inherited and acquired deficits in mitochondrial function act upstream of classical AD pathology and drive progressive network failure^12^. Furthermore, our recent systems-level analyses of asymptomatic AD (AsymAD), individuals who remain cognitively intact despite substantial amyloid and tau pathology, identified preserved mitochondrial bioenergetics as a hallmark of resilience^13^.

Relative to symptomatic AD, AsymAD brains maintained oxidative phosphorylation, electron transport chain activity, fatty acid oxidation, and branched-chain amino acid metabolism, indicating sustained metabolic flexibility and energy homeostasis. These observations support the emerging concept that mitochondrial dysfunction is a critical determinant of disease progression and suggest that therapeutic restoration of mitochondrial function may confer benefit even in the presence of established AD pathology^13^.

Despite decades of research, disease-modifying therapies for AD remain limited. Anti-amyloid monoclonal antibodies provide only modest clinical benefit while carrying significant risks and do not address the underlying metabolic dysfunction that emerges early in the disease. Emerging evidence supports that amyloid accumulation is itself a downstream consequence of metabolic dysfunction, as impaired mitochondrial metabolism and bioenergetic failure have been shown to precede and drive amyloid pathology in AD^2,14^. These observations have renewed interest in therapeutic strategies that restore metabolic resilience rather than targeting individual pathological endpoints. We previously demonstrated that weak inhibition of mtCI with the brain-penetrant small molecule CP2 activates an adaptive stress response that restores mitochondrial function, improves glucose metabolism, enhances synaptic activity, reduces inflammation and oxidative stress, and improves cognition in symptomatic APP/PS1 mice^15^. Chronic treatment initiated after the onset of neurodegeneration produced sustained efficacy over 14 months, demonstrating that modulation of a single mitochondrial target can engage a broad neuroprotective program^15^. Recent studies further indicate that restoring mitochondrial function alone is sufficient to improve cognitive function in mice^16^. However, whether this approach remains effective during advanced stages of disease, when metabolic dysfunction and network collapse are already established, remains unknown.

Here, we address this critical question by examining the effects of CP2 treatment initiated at 19 months of age in APP/PS1 female mice. To define the metabolic mechanisms underlying therapeutic response, we developed *iMiceBrain*, the first brain-specific genome-scale metabolic model of the mouse brain, enabling systems-level simulation of disease-associated metabolic rewiring. By integrating transcriptomics, targeted metabolomics, lipidomics, and genome-scale metabolic modeling, we demonstrate that mild mtCI modulation reprograms cholesterol and lipid metabolism, enhances mitochondrial substrate flexibility, restores metabolic resilience, and reverses AD-associated molecular signatures even at advanced stages of disease. These findings identify restoration of mitochondrial metabolic resilience as a disease-modifying strategy and establish mtCI modulation as a therapeutic approach capable of re-engaging protective biological programs in the aging AD brain.

## Results

### APP/PS1 female mice recapitulate the metabolic, inflammatory, and transcriptional signatures of human AD

To determine whether mtCI modulation is efficacious in older AD mice, we assemble treatment cohorts of female APP/PS1 mice linked to familial AD (FAD)^17^, and non-transgenic (NTG) age-and sex-matched littermates. Separate cohorts were treated with CP2 or vehicle for 6 weeks starting at 19 months of age. The APP/PS1 model was selected in part because its slower disease progression, relative to more aggressive models such as the 5xFAD, allows aging-related pathological processes to develop alongside amyloid deposition. While this model relies on familial AD-linked mutations and does not fully recapitulate the etiology of LOAD, studying it at an advanced age (20-21 months) may better capture the interplay between amyloid pathology and age-dependent cellular dysfunction that characterizes the human disease. To characterize the transcriptome landscape of late-stage AD, we profiled APP/PS1 and NTG littermates after 6 weeks of treatment (*n* = 5/group, **Fig. 1a**). Brain hemispheres were processed separately: the left hemisphere was used for bulk RNA sequencing, targeted metabolomics, and untargeted lipidomics, whereas the right hemisphere was used for Western blot (WB) analyses (**Fig. 1a**).

**Figure 1:**
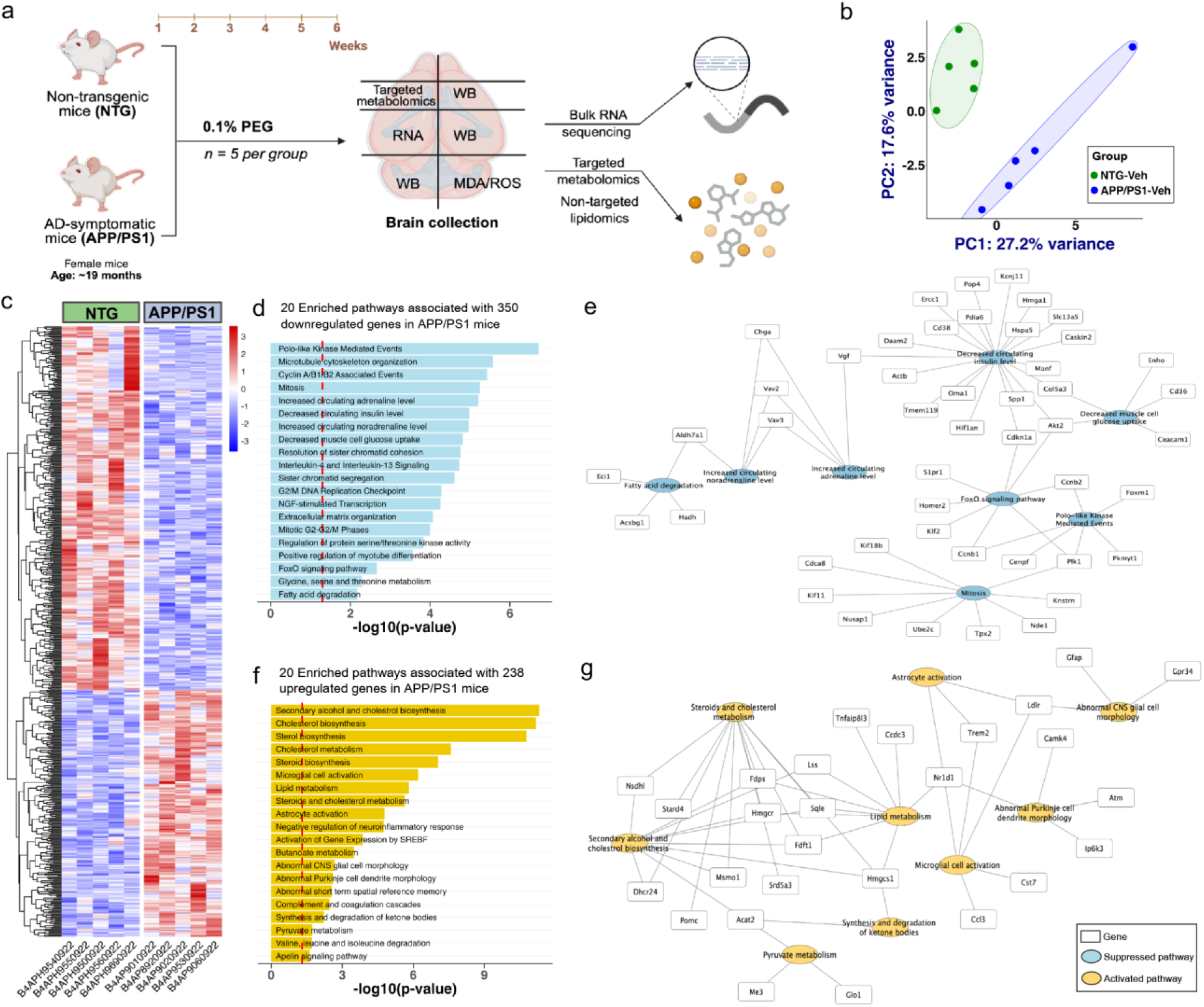
APP/PS1 mice exhibit molecular changes associated with AD. (a) Study design: Brain tissues from NTG and APP/PS1 mice were used for transcriptomics, metabolomics, and lipidomics analyses. (b) PCA of normalized gene expression data from mouse brain tissue. (c) Heatmap visualizing the normalized gene expression data of differentially expressed genes (DEGs) identified in APP/PS1 mice. (d) Enriched biological pathways associated with downregulated genes identified in APP/PS1 mice. The statistical significance of enrichment is indicated by the –log10 *p*-value; a vertical, red line marks a threshold of 1.3, corresponding to a *p*-value of 0.05. (e) Network analysis indicating gene-pathway interaction of downregulated genes. Blue nodes represent downregulated pathways associated with the genes in rectangular boxes. (f) Enriched biological pathways associated with upregulated genes identified in APP/PS1 mice. The statistical significance of enrichment is indicated by the –log10 *p*-value; a vertical, red line marks a threshold of 1.3, corresponding to a *p*-value of 0.05. (g) Network analysis indicating gene-pathway interaction of upregulated genes. Yellow nodes represent upregulated pathways associated with the genes in rectangular boxes. *Created in BioRender. Gabal, E. (2026)* https://BioRender.com/2kdkf8r.

Differential gene expression analysis was restricted to protein-coding genes (absolute log2-fold change (|log_2_FC|) ≥ 0.25, *p* < 0.01; **Supplementary Table 1**). Principal component analysis (PCA) showed distinct transcriptomic separation between APP/PS1 and NTG mice (PC1: 27.2%, PC2: 17.6% variance; **Fig. 1b**). Differential gene expression analysis identified 350 downregulated and 238 upregulated genes in APP/PS1 vs NTG mice (**Fig. 1c**). Gene set enrichment analysis (GSEA) of downregulated genes showed suppression of mitotic and cell cycle pathways as the dominant transcriptomic signature in APP/PS1 mice, with Polo-like kinase-mediated events, Cyclin A/B1/B2-associated events, G2/M DNA replication checkpoint, and sister chromatid segregation among the most significantly enriched terms (*Ccnb1/2*, *Foxm1, Cenpf,* and *Pkmyt1)* (**Fig. 1d**). FoxO signaling was similarly suppressed (*Homer2*, *S1pr1*, *Akt2*, *Klf2*, and *Cdkn1a)* in APP/PS1 mice. Network visualization indicated mitosis as a central hub, with genes including *Plk1, Foxm1, Ccnb1/2, Cenpa,* and *Cenpf* converging on mitotic regulation (**Fig. 1e**). Beyond cell cycle suppression, downregulated genes were associated with decreased circulating insulin levels, reduced muscle cell glucose uptake, and FoxO signaling, indicating impaired metabolic sensing and insulin signaling, along with suppression of fatty acid degradation and extracellular matrix organization (**Fig. 1e**).

Upregulated genes in APP/PS1 mice were enriched for sterol and cholesterol biosynthesis pathways, with secondary alcohol and cholesterol biosynthesis, and SREBF-driven gene activation representing the most significant terms (**Fig. 1f**). Network analysis linked genes including *Hmgcr, Hmgcs1, Lss, Sqle, Fdps, Dhcr24, Nsdhl, Msmo1*, and *Acat2* to coordinated dysregulation of the mevalonate pathway and lipid metabolism (**Fig. 1g**). Microglial cell activation and astrocyte activation emerged as prominent upregulated pathways, with *Trem2, Cst7, Ccl3, Gpr34, Gfap*, and *Ldlr* forming a neuroinflammatory gene network associated with abnormal CNS glial morphology (**Fig. 1f, g**). Synthesis and degradation of ketone bodies and pyruvate metabolism were also upregulated, suggesting compensatory shifts in energy substrate utilization. This analysis highlights the transcriptomic landscape in late-stage APP/PS1 mice defined by mitotic failure, insulin resistance, aberrant cholesterol biosynthesis, and neuroinflammation, mirroring key metabolic and inflammatory features of human AD.

### CP2 treatment modulates key signaling pathways associated with neuroprotection in aged APP/PS1 mice

To determine whether CP2 retains efficacy at advanced disease stages, we performed differential gene expression analysis on brain tissue from 20-21-month-old APP/PS1 and NTG female mice treated with CP2 or vehicle for 6 weeks, accounting for the genotype-by-treatment interaction. PCA indicated APP/PS1 mice clustering distinctly from NTG group (**Fig. 2a**). CP2 treatment in APP/PS1 mice resulted in 113 downregulated and 59 upregulated genes (|log₂FC| ≥ 0.25, *p*<0.01; **Fig. 2b**), representing a more focused transcriptional response compared to the broad reprogramming observed in vehicle-treated APP/PS1 mice.

**Figure 2.**
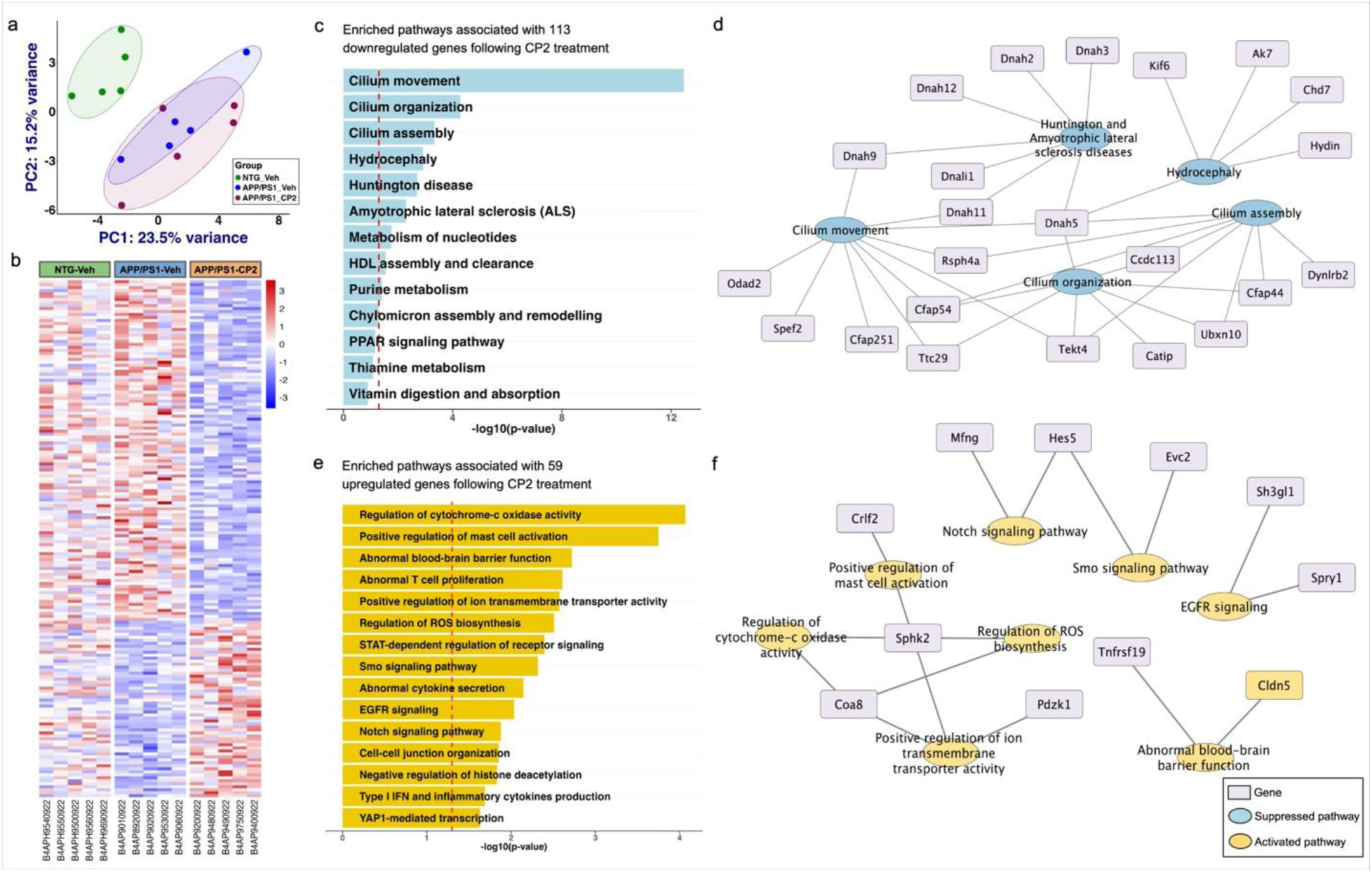
CP2 treatment enhances targeted neuroprotective transcriptional program in APP/PS1 female mice. (a) Principal component analysis (PCA) of NTG, APP/PS1 vehicle and APP/PS1 CP2 treated groups. (b) Heatmap visualizing the normalized gene expression data of DEGs following CP2 treatment (113 downregulated and 59 upregulated genes) for the three experimental groups (NTG, APP/PS1 vehicle, APP/PS1 CP2-treated; *n* = 5/group). (c) Enriched biological pathways associated with downregulated genes identified in CP2-treated APP/PS1 mice. The statistical significance of enrichment is indicated by the –log10 *p*-value, with the vertical, red line representing a threshold of 1.3, corresponding to a *p*-value of 0.05. (d) Network analysis indicating gene-pathway interaction of downregulated genes. Blue nodes represent downregulated pathways associated with the genes in rectangular boxes. (e) Enriched biological pathways associated with upregulated genes identified in CP2-treated APP/PS1 mice. The statistical significance of enrichment is indicated by the –log10 *p*-value, with the vertical, red line representing a threshold of 1.3, corresponding to a *p*-value of 0.05. (f) Network analysis indicating gene-pathway interaction of upregulated genes. Red nodes represent upregulated pathways associated with the genes in rectangular boxes. *Created in BioRender. Gabal, E. (2026)* https://BioRender.com/o0b9zlm.

Genes downregulated by CP2 treatment in APP/PS1 mice were enriched in cilia morphology and structure, neurodegenerative disease pathways, and nucleotide metabolism (**Fig. 2c**). Network analysis showed cilia structure and motility genes, including *Dnah5, Dnah9, Dnah11, Rsph4a, Cfap54,* and *Hydin* as central nodes within these suppressed pathways (**Fig. 2d**). *Dnali1* has been implicated in neurodegeneration as a suppressor of autophagy^18^. Reduced expression of *Chd7, Hydin, Ak7*, and *Kif6* (**Fig. 2c, d**), implicated in neurodevelopment and CSF circulation^19^. This suggests that CP2 may improve autophagy, CSF homeostasis and ventricular dynamics, processes linked to impaired Aβ clearance in AD.

Upregulated genes were enriched in mitochondrial function, neuroinflammatory modulation, blood-brain barrier (BBB) integrity, and ciliary signaling, including *Smo*, *Notch*, and *EGFR* pathways, epigenetic regulation linked to neuronal repair and homeostasis (**Fig. 2e**). *Sphk2,* encoding sphingosine kinase 2, featured in top four upregulated pathways, indicating sphingosine-1-phosphate (S1P) signaling in CP2-mediated neuroinflammatory modulation (**Fig. 2e, f**). *Smo* signaling pathways enrichment was driven primarily by *Hes5* and *Evc2* (**Fig. 2e, f**). Upregulation of *Cldn5 (*Claudin-5) and *Tnfrsf19* indicated maintenance of BBB integrity (**Fig. 2e, f**)^20^. Our analysis indicates CP2 treatment elicits a targeted neuroprotective transcriptional program that counteracts AD-associated molecular disturbances and promotes restoration of neuronal and vascular homeostasis.

### CP2 reverses AD-associated molecular changes by enhancing neuroprotection and proteostasis

Comparative analysis of genotype- and CP2-regulated genes identified 23 shared DEGs, of which 20 were downregulated in APP/PS1 mice but restored by CP2 treatment (**Fig. 3a, Supplementary Table 2**). These genes are involved in various biological processes, including neurogenesis, neuroprotection, lipid metabolism, and proteostasis. Among them, circadian clock components *Dpb* and *Ciart* were downregulated in APP/PS1 mice, and the clock-controlled gene *Cldn5* was restored to NTG-comparable levels following CP2 treatment (**Fig. 3b**). In APP/PS1 mice, CP2 treatment resulted in a coordinated decrease in *Ahcy* and increased *Ahcyl* expression, normalizing both to NTG levels and suggesting restoration of circadian transcriptional activity (**Fig. 3b**). Surprisingly, APP/PS1 mice exhibited a near-complete loss of *Neurog1* and *Nfe2*, key regulators of neurogenesis and neuroprotection, respectively, which CP2 treatment restored to NTG levels (**Fig. 3b**). These findings underscore CP2’s capacity to support neuroprotective programs in aged AD brain.

**Figure 3.**
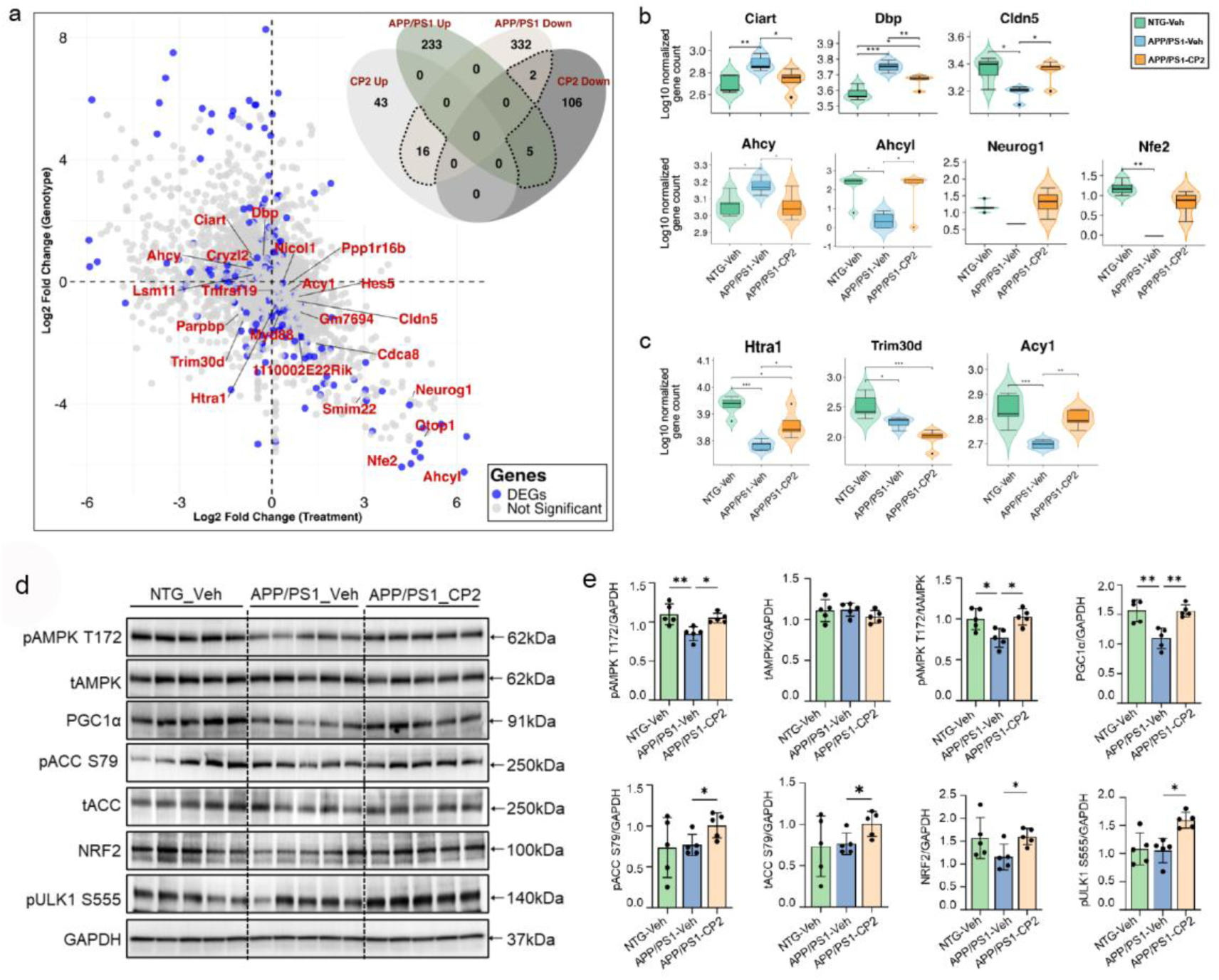
CP2 treatment activates the mechanism of action in the brain of aged APP/PS1 mice, enhancing neuroprotection and proteostasis. (a) The genotype-by-treatment plot showing 23 differentially expressed genes from the comparison of the APP/PS1 genotype and the CP2 treatment. The genes are derived from the dotted portion in the inset Venn diagram. (b) Violin plot of genes associated with circadian rhythm, neuroprotection and neurogenesis functions. (c) Violin plot of genes associated with proteostasis function. Statistical significance was determined using one-way ANOVA followed by Tukey’s HSD post-hoc test (*p* < 0.05). (d) Western blot analysis of brain tissue confirmed activation of AMPK and the downstream neuroprotective mechanisms in aged APP/PS1 mice. Each lane represents a mouse, *n* = 5 mice *per* group. Statistical analysis was performed using ordinary one-way ANOVA followed by Bonferroni post hoc tests for multiple comparisons between APP/PS1 Vehicle, NTG Vehicle, and APP/PS1 CP2 groups. \**P* < 0.05; \*\**P* < 0.01. *Created in BioRender. Gabal, E. (2026)* https://BioRender.com/3e8swj3.

CP2 treatment also targeted core regulators of proteostasis, a process suppressed by both aging and AD^21^. We found that CP2 treatment increased the expression of key regulators that mediate autophagy and proteome homeostasis (**Fig. 3c, d**). *Htra1*, a protease and chaperone involved in misfolded protein clearance, was downregulated in APP/PS1 mice and restored by CP2 treatment. CP2 also upregulated *Acy1*, which maintains protein metabolism through amino acid recycling, and *Tmem258*, a component of the oligosaccharyltransferase complex with a regulatory role in ER homeostasis (**Fig. 3c**).

We next determined whether CP2 engages its intended mechanism of action and activates downstream neuroprotective signaling pathways in the aged AD brain^3,22–25^. Western blot analyses confirmed activation of the AMPK signaling cascade and its downstream targets in brain tissue from CP2-treated APP/PS1 mice (**Fig. 3d, Supplementary Fig. 1a**). Levels of pAMPK, a central regulator of cellular energy homeostasis, as well as the mitochondrial biogenesis regulator peroxisome proliferator-activated receptor-γ coactivator-1α (PGC1α) and nuclear factor E2-related factor 2 (NRF2), were significantly reduced in vehicle-treated APP/PS1 mice compared with NTG littermates, indicating impaired AMPK-PGC1α-NRF2 signaling and antioxidant defenses (**Fig. 3d,e, Supplementary Fig. 1b**). Consistent with previous reports, CP2 treatment activated AMPKα and its downstream target acetyl-CoA carboxylase (ACC), confirming target engagement. Importantly, CP2 restored the expression of pAMPK, PGC1α, and NRF2 to levels comparable to those observed in NTG mice. In agreement with AMPK activation, CP2 also increased phosphorylation of Unc-51-like autophagy activating kinase 1 (ULK1) at Ser555, indicative of enhanced autophagy (**Fig. 3e**, **Supplementary Fig. 1c**). These findings demonstrate that CP2 effectively engages its mechanism of action in the aged brain and restores key neuroprotective, mitochondrial, and proteostatic signaling pathways that are compromised in APP/PS1 mice.

### *iMiceBrain*: the first mouse brain-specific genome-scale metabolic reconstruction with translational relevance to human brain metabolism

To resolve the metabolic flux changes underlying CP2’s transcriptional reprogramming, we turned to constraint-based modeling, which integrates gene expression data with stoichiometric and thermodynamic constraints to simulate how metabolites flow through biochemical networks^26^. For this purpose, we developed *iMiceBrain*, the first mouse brain genome-scale metabolic reconstruction (GEM), constructed as a consensus model using four pruning algorithms (**Fig. 4a**). The resulting network comprises 6042 reactions, 3111 metabolites, and 1668 metabolic genes spanning 89 metabolic subsystems, with transport, lipid, and energy metabolism most prominently represented (**Fig. 4b**). Brain specificity was confirmed by cross-referencing *iMiceBrain* genes against mouse brain transcriptomic data from the Human Protein Atlas (HPA)^27^ and Allen Institute^28^, yielding 94% and 83.3% gene-set overlap, respectively (**Fig. 4c**). To assess translational relevance, human homologs of *iMiceBrain* genes were mapped via HomoloGene and compared against Recon3D^29^ and the human brain GEM^30^. The vast majority of *iMiceBrain* genes had identifiable human homologs, with only 13 genes (∼0.8%) lacking homology (**Fig. 4c; Supplementary Table 3**). Notably, *iMiceBrain* uniquely contains five human homologous genes, namely *Slc22a25, Prodh, Slc22a19, Uqcrh*, and *Slc25a31* that are absent from both existing human models, highlighting the model’s capacity to capture brain metabolic features not represented in current human GEMs (**Fig. 4d**).

**Figure 4.**
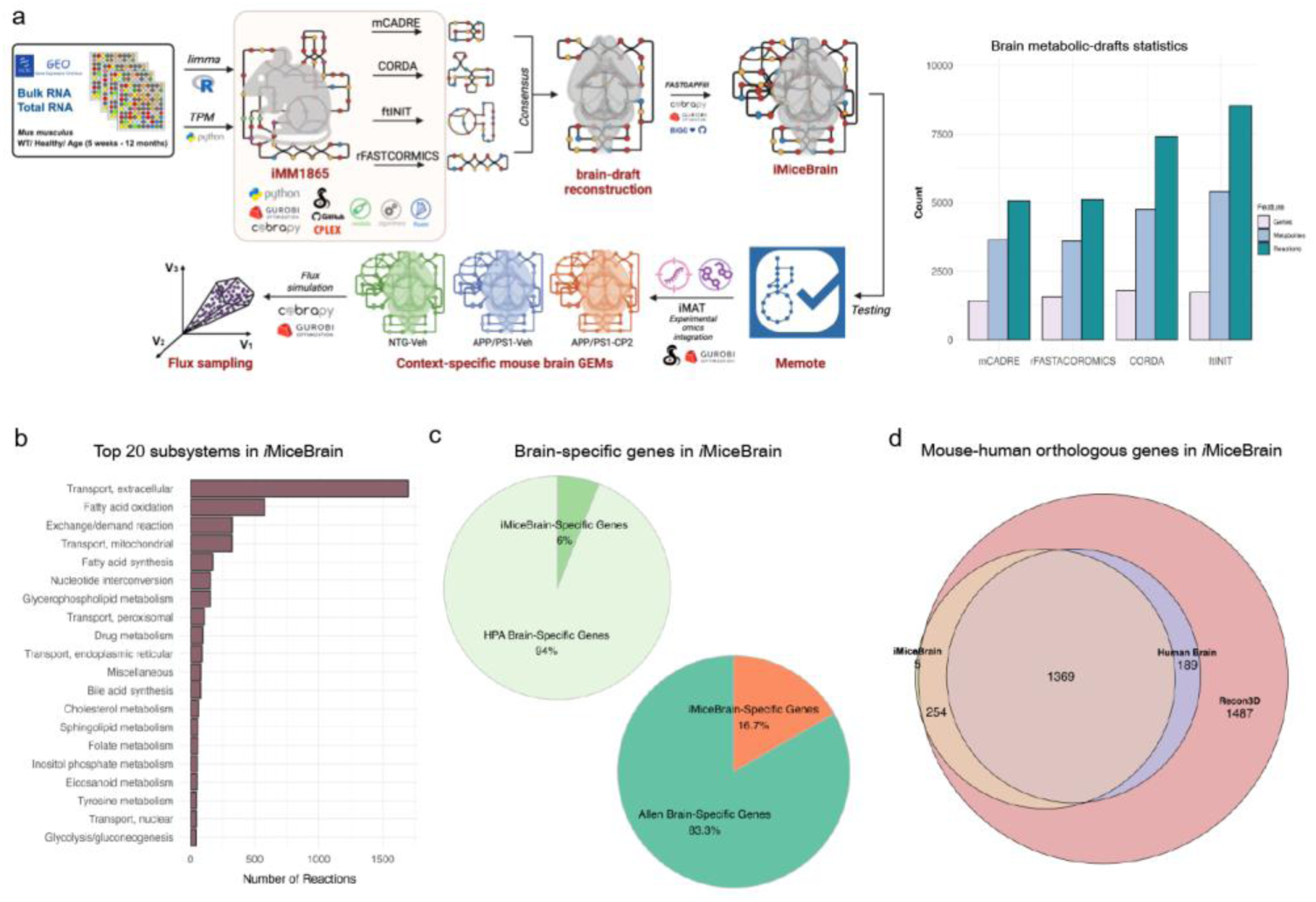
Details of genome-scale metabolic reconstruction of mouse brain, *iMiceBrain*. (a) The genome-scale metabolic reconstruction of the mouse brain was generated using four pruning algorithms of mCADRE, rFASTACOROMICS, CORDA, and ftINIT, from which the consensus brain model was developed. The number of reactions, metabolites, and genes in the draft reconstructions is indicated in bar graphs. (b) Top 20 metabolic subsystems in *iMiceBrain* corresponding to the number of reactions in the network. (c) Pie charts representing *i*MiceBrain genes mapped to mouse brain transcriptomics data acquired from Human Proteome Atlas (HPA) and Allen Institute. (d) Comparison of the human-mouse orthologous genes present in *iMiceBrain* versus genome-scale metabolic network of human (Recon3D) and the human brain model (from whole body models). *Created in BioRender. Gabal, E. (2026)* https://BioRender.com/xnazjnw.

Model quality was assessed using Memote^31^, a standardized benchmarking tool that scores genome-scale metabolic reconstructions across structural integrity, stoichiometric consistency, metabolite connectivity and annotation completeness. *iMiceBrain* received an overall score of 88%. The metabolic reconstruction exhibited no orphan metabolites, blocked reactions, or dead-end metabolites, with full stoichiometric consistency and metabolite connectivity (100%; **Supplementary File 1**). Functionality testing against universally required mammalian metabolic tasks confirmed performance of 80% of essential tasks, with flux consistency validated and confirmed ATP production from glucose under both aerobic and anaerobic conditions (**Supplementary Table 4**). As a standard validation of GEM predictive accuracy^32,33^, single gene knockout simulation results were compared against experimentally validated lethal and non-lethal genes (lethality threshold: ≥30% reduction in predicted growth). The results showed 100% sensitivity for viable genes (107/107) and correctly predicted lethality for 7 of 55 lethal genes (12% sensitivity), outperforming the parental model *i*MM1865 (5%), likely reflecting the exclusion of non-brain alternative reactions in the tissue-specific reconstruction (**Supplementary Table 5**). Collectively, *iMiceBrain* is a robust, biologically consistent metabolic reconstruction that accurately captures core brain metabolic processes and offers a powerful platform for systems-level investigation of neurological disease and therapeutic response.

### *iMiceBrain* identifies CP2-mediated reprogramming of lipid and cholesterol metabolism

To mechanistically resolve CP2’s effects on brain metabolism, we integrated brain transcriptomics with *iMiceBrain* using iMAT, generating 15 condition-specific metabolic reconstructions representing *in silico* metabolic profiles across all experimental groups. Monte Carlo flux sampling followed by Kolmogorov-Smirnov testing was applied to characterize metabolic flux distributions across conditions. *iMiceBrain* simulations identified CP2-driven remodeling of cholesterol biosynthesis as a primary metabolic shift. Reactions mediated by HMGCOAR, HMGCOARx, and MEVK1, the initiating steps of the mevalonate pathway, showed broad flux distributions in NTG and CP2-treated APP/PS1 mice but were largely constrained near zero in vehicle-treated APP/PS1 mice (**Fig. 5a**), indicating mevalonate pathway suppression in disease that is restored by CP2 treatment. Consistent with this, transcriptomic analysis of APP/PS1 CP2-treated mice identified significant suppression of key cholesterol biosynthesis genes, including *Hmgcs1, Hmgcr, Fdft1*, and *Lss* (**Fig. 5b,c**). There was downregulation of lipid transport genes *Lcat* and *Apoa1*, which mediate cholesterol efflux and HDL remodeling (**Fig. 5b,d**; **Supplementary Table 6**). These findings indicate that mechanistically, CP2 recalibrates cholesterol biosynthesis and transport to restore lipid equilibrium in the aged AD brain.

**Figure 5.**
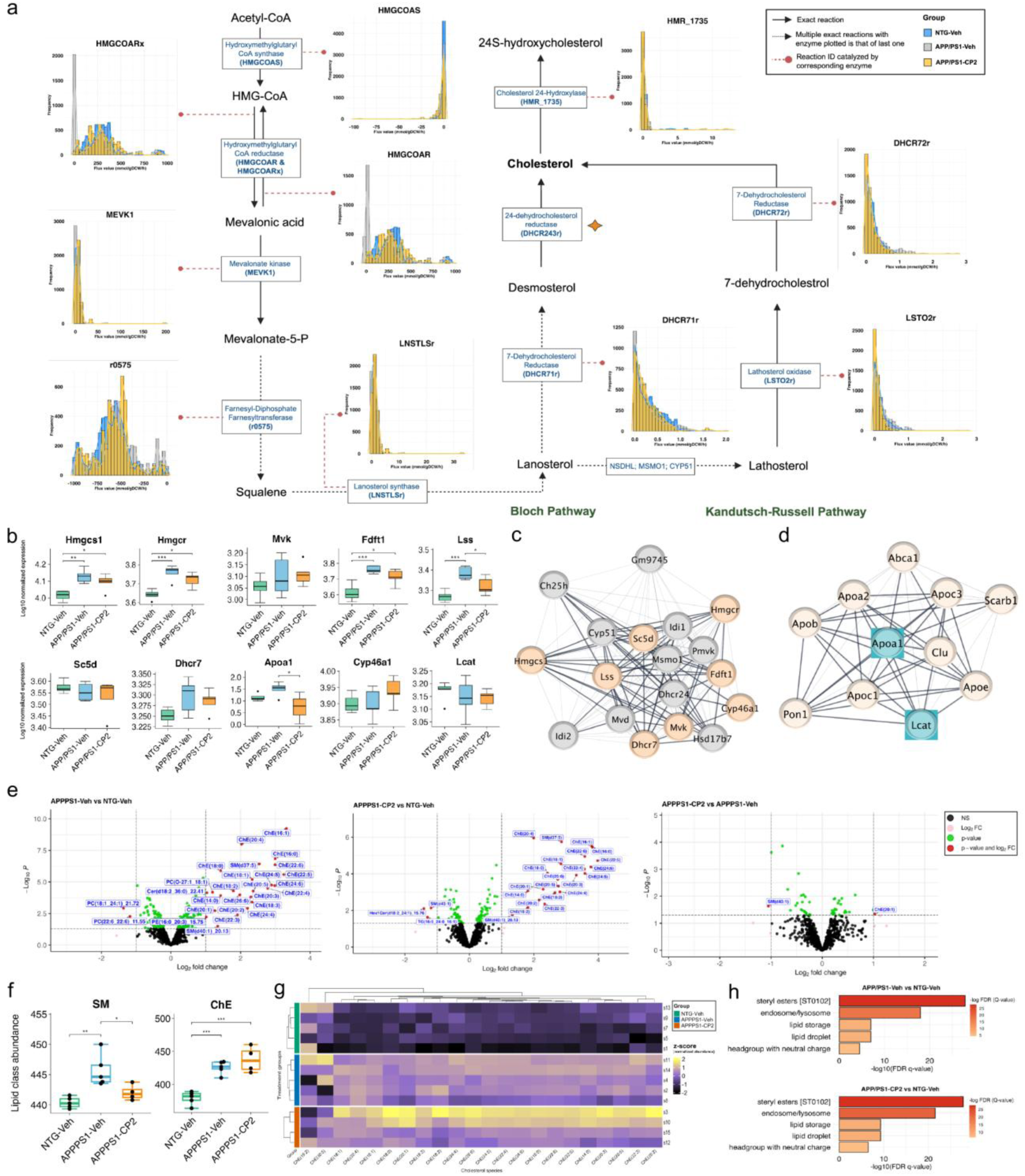
Metabolic simulation and omics analysis elucidate the influence of CP2 on cholesterol metabolism reprogramming. (a) Flux distributions of key reactions involved in cholesterol biosynthesis across all context-specific models. Histograms represent distributions obtained via Monte Carlo flux sampling (thinning factor of 1000). (b) Box plots showing normalized expression of genes involved in cholesterol biosynthesis and clearance across groups. (c) STRING protein-protein interaction (PPI) network of cholesterol biosynthesis proteins in the Bloch pathway, identified from the analysis (salmon-colored) and their interacting partners. (d) STRING PPI network of cholesterol clearance proteins in the Kandutsch-Russell pathway, identified from the analysis (cyan-colored) and their interacting partners. Bold edges in STRING maps denote high-confidence interactions (interaction score > 0.9). (e) Volcano plots of differentially abundant lipid species identified from lipidomics analysis following |log2FC| > 1.0 and adjusted *p*-value < 0.05 cut-off. (f) Boxplot representing levels of sphingomyelin (SM) and cholesteryl ester (ChE) species. (g) Heatmap representing normalized levels of cholesteryl esters identified from lipidomics analysis. (h) Enriched pathways of differential lipid species using LION database. Statistical significance was determined via one-way ANOVA followed by Tukey’s HSD post-hoc test (*p* < 0.05). *Created in BioRender. Gabal, E. (2026)* https://BioRender.com/iakskk9.

Lipidomic profiling of mouse brain tissue corroborated *iMiceBrain* predictions, revealing cholesteryl esters (ChEs) as the most abundant lipid class across groups, followed by phosphatidylcholine (PC) and sphingomyelin (SM) species. ChEs were elevated in APP/PS1 mice relative to NTG controls and their levels further increased by CP2 treatment (**Fig. 5e-g**), consistent with enhanced lipid droplet-mediated buffering of excess cholesterol to limit lipotoxicity. This mechanism is mediated by CP2 to restore lipid homeostasis. Steryl esters, neutral lipids (triacylglycerols, diacylglycerols, and phosphatidylcholines), and lipid storage pathways were the most significantly enriched by CP2 treatment (**Fig. 5h**). PC species such as PC(18:1/24:1) and PC(22:6/22:6), which are critical for membrane integrity and synaptic vesicle function^34^, were reduced in APP/PS1 mice (**Fig. 5e**), suggesting structural and synaptic membrane disruption consistent with the transcriptomic findings.

CP2 treatment significantly reduced total sphingomyelin (SM) levels in APP/PS1 mice (**Fig. 5f**). Sphingolipid species such as SM(d43:1) and Hex1Cer(d18:2/24:1) are associated with pro-inflammatory signaling, insulin resistance, and elevated AD risk^35^. This sphingolipid remodeling in CP2-treated APP/PS1 mice suggests a shift away from neuroinflammatory lipid signaling toward neuroprotective membrane composition. Additionally, CP2 reduced triglyceride (TG) levels, suggesting enhanced utilization of triglyceride-derived fatty acids for mitochondrial β-oxidation and improved bioenergetic capacity. Integration of *in silico* flux analysis with multi-omics data using *iMiceBrain* demonstrates that CP2 restores cholesterol biosynthesis, lipid storage, sphingolipid signaling, and energy substrate utilization, thereby facilitating metabolic homeostasis in the aged AD brain.

### CP2 treatment enhances mitochondrial bioenergetic capacity through β-oxidation and pyruvate flux reprogramming

*iMiceBrain* flux sampling identified CP2-specific activation of the carnitine-acylcarnitine translocase reaction (ACRNTm), which mediates the import of long-chain fatty acids into mitochondria for β-oxidation^36^. The flux for this reaction is absent in both NTG and APP/PS1 mice treated with vehicle (**Fig. 6a**). Citrate synthase flux (CSm) was reduced in CP2-treated APP/PS1 mice compared to NTG and APP/PS1 groups, while pyruvate oxidation to acetyl-CoA (PDHm) was detected exclusively in vehicle- and CP2-treated APP/PS1 mice (**Fig. 6a**). Together, these simulations support a CP2-induced shift in substrate utilization toward fatty acids and altered partitioning of acetyl-CoA. Lipidomic and metabolomic profiling corroborated these predictions. Although not statistically significant, brain lipidomics showed trends toward reduced triglycerides and elevated fatty acids and acylcarnitines following CP2 treatment (**Fig. 6b**), consistent with increased fatty acid utilization. Targeted metabolomics identified acylcarnitines as key discriminators across groups by orthogonal partial least-squares-discriminant analysis (oPLS-DA) (**Fig. 6c**), with trends toward increased acetyl-CoA and reduced pyruvate levels in CP2-treated APP/PS1 mice (**Fig. 6d, Supplementary Fig. 2**). This result is consistent with enhanced pyruvate dehydrogenase (PDH)-dependent pyruvate-to-acetyl-CoA conversion.

**Figure 6.**
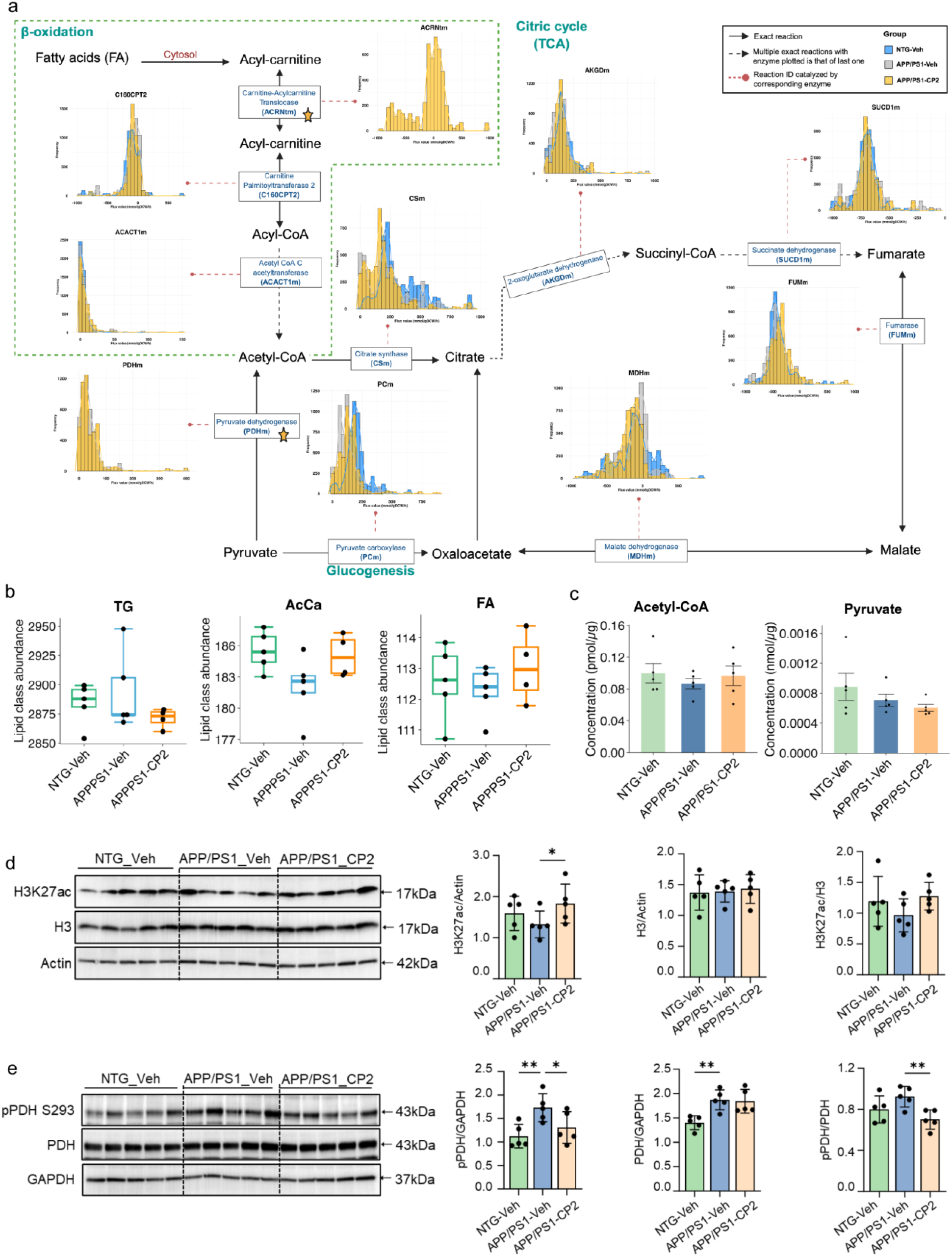
CP2 treatment improves acylcarnitine-mitochondrial metabolism in APP/PS1 mice. (a) Flux distribution of metabolic reactions involved in β-oxidation, the TCA cycle, and glycogenesis, associated with mitochondrial function. Histograms represent flux distributions calculated using Monte Carlo flux sampling (thinning factor of 1000). (b) Boxplots representing the total abundance of lipid species in the following classes: triglycerides (TG), acylcarnitine (AcCa), and fatty acids (FA). (c) Acetyl-CoA and pyruvate concentrations were identified by targeted metabolomics. (d and e) Western blot analysis of brain tissue from APP/PS1 mice treated with CP2 for 6 weeks shows increased H3K27ac and reduced phosphorylation of pyruvate dehydrogenase (pPDH). Each lane represents an individual mouse, and *n = 5* mice per group. Statistical significance was determined via the Kruskal-Wallis test and one-way ANOVA followed by Fisher’s LSD post-hoc test. *P < 0.05, **P < 0.01. *Created in BioRender. Gabal, E. (2026)* https://BioRender.com/nc1rf8e.

To assess whether changes in acetyl-CoA availability influence chromatin state, we examined histone H3 lysine 27 acetylation (H3K27ac), an active enhancer-associated histone mark sensitive to acetyl-CoA levels. Western blot analysis showed reduced H3K27ac in vehicle-treated APP/PS1 mice, which was restored by CP2 treatment (**Fig. 6e, Supplementary Fig. 3**). A similar effect was observed in CP2-treated human SH-SY5Y cells expressing APP with the Swedish mutation (data not shown), suggesting a conserved epigenetic response. Consistent with increased PDH activity, CP2-treated APP/PS1 mice exhibited a reduced pPDH/PDH ratio (**Fig. 6e**). Integrating flux simulations with lipidomic and metabolomic data, these findings indicate that CP2 promotes coordinated engagement of fatty acid utilization and PDH-dependent pyruvate metabolism, supporting flexible substrate use, maintenance of TCA cycle function, and increased acetyl-CoA availability for bioenergetic and epigenetic processes under chronic metabolic stress in the aged AD brain.

## Discussion

AD is a multifactorial neurodegenerative disease driven by convergent disruptions in energy and lipid homeostasis, proteostasis, mitochondrial function, and glial reactivity^2^. We previously demonstrated that weak inhibition of mtCI with the small molecule CP2 activates an adaptive stress response that preserves synaptic function and blocks ongoing neurodegeneration across multiple AD mouse models, engaging AMPK-dependent signaling, autophagy, and mitochondrial biogenesis to reduce pTau, Aβ, oxidative stress, and inflammation while restoring energy homeostasis in brain and periphery^3,15,22,23^. Cross-validation with human RNA-seq data confirmed that CP2 positively modulates disease-relevant pathways, including inflammation, oxidative stress, and synaptic function^15,22,23^. Here, using a systems-level approach integrating transcriptomics, lipidomics, metabolomics, and the novel brain-specific metabolic model *iMiceBrain*, we demonstrate that CP2 drives broad metabolic reprogramming even in late-stage AD, restoring neuroprotective, proteostatic, and bioenergetic programs in APP/PS1 mice.

Transcriptomic profiling of 20-21-month-old APP/PS1 mice revealed widespread molecular disruption consistent with advanced AD. Suppression of mitotic and cell cycle genes indicates impaired proliferative and DNA repair capacity, while downregulation of FoxO and PPAR signaling pathways reflects perturbations in insulin signaling, lipid metabolism, and energy sensing. These findings are consistent with the established link between impaired brain insulin signaling and the mechanisms studied in AD. Aberrant FoxO activation through disrupted insulin signaling may drive maladaptive transcriptional programs, compounding lipid and amino acid metabolic dysregulation^37^. Prominently upregulated cholesterol biosynthesis genes align with prior reports of cholesterol accumulation in Aβ plaques in both AD mouse models and patients ^38^, while concurrent activation of microglial and astrocytic markers reflects the neuroinflammatory signatures characteristic of AD, where reactive glia concentrate around amyloid deposits and exacerbate disease progression^39^.

Our data suggest that even at the advanced disease stage, CP2 elicited a targeted neuroprotective transcriptional response in APP/PS1 mice. These effects can be understood within the broader context of astrocyte-neuron metabolic coupling, which is increasingly recognized as an upstream vulnerability in AD. Neurons depend on the astrocyte-neuron lactate shuttle (ANLS) for metabolic support, with astrocytic glycolysis and glycogenolysis providing lactate required for long-term potentiation, synaptic plasticity, and memory consolidation^40^. Disruption of this shuttle through indoleamine-2,3-dioxygenase 1 (IDO1)-mediated suppression of astrocytic glycolysis produces cognitive and synaptic deficits upstream of amyloid and tau pathways^14^, positioning ANLS failure as a cause rather than a consequence of classical AD pathology. This metabolic vulnerability is compounded by the cholesterol burden imposed on glia during neurodegeneration: neurons transfer activity-induced toxic fatty acids to astrocytes via ApoE-containing lipid particles for storage in lipid droplets and subsequent mitochondrial β-oxidation, a pathway that fails when astrocytic mitochondria are compromised, leading to pathological lipid accumulation and neuroinflammation^41^. In human iPSC-derived astrocytes, excess cholesterol is sequestered in lipid droplets that fail to form productive contacts with mitochondria, preventing oxysterol production and downstream LXRα-mediated signaling required for APOE lipoprotein lipidation and cholesterol efflux, while lipid-laden microglia lose their capacity for neuronal surveillance and debris clearance^42^. Our finding that CP2 simultaneously restores mitochondrial β-oxidation capacity, activates the carnitine-acylcarnitine shuttle, normalizes cholesterol biosynthesis gene expression, and enhances cholesterol ester buffering suggests that CP2 re-enables precisely the mitochondria-dependent lipid processing axis that becomes overwhelmed in AD. This is suggestive of restoring astrocytic oxidative capacity to handle cholesterol-rich debris while re-establishing the bioenergetic foundation upon which the ANLS depends. Critically, AMPK, which CP2 activates, directly regulates the astrocytic lactate shuttle and is required for non-cell-autonomous neuronal survival^43^, suggesting that CP2’s target engagement of mtCI may restore metabolic coupling between astrocytes and neurons by expanding both the lipid-handling and energy-supply arms of astrocytic support simultaneously.

CP2 upregulated *Sphk2*, consistent with restoration of S1P metabolism that is impaired by Aβ deposition in post-mortem AD brains, alongside Notch signaling components critical for cerebrovascular and neural integrity^44^. CP2 also restored circadian clock gene expression (*Dbp, Ciart, Cldn5*), addressing a system disrupted in approximately 80% of AD patients, and rescued near-complete loss of *Neurog1*, a proneural transcription factor whose enhanced expression has been associated with slower cognitive decline and neurodegeneration in cognitively impaired individuals^45^. CP2 restored proteostatic balance by upregulating *Htra1*, a protease implicated in preventing α-synuclein aggregation and degrading amyloid fibrils and identified as a candidate gene in human AD (AMP-AD)^46^, while suppressing the proteasome assembly inhibitor *Psmd5* and restoring aminoacylase-1 expression, whose deficiency has been linked to neurological and psychiatric deficits.

The CP2-driven upregulation of PGC1α and NRF2 may represent a mechanistic node linking mitochondrial restoration to the lipid metabolic remodeling observed in this study. PGC1α drives mitochondrial biogenesis to expand oxidative capacity, while NRF2 coordinates the antioxidant defense necessary to sustain β-oxidation without excessive ROS generation. Together, these programs re-enable the mitochondria-dependent lipid processing that is essential for astrocytic function: neurons transfer activity-induced toxic fatty acids to astrocytes for lipid droplet storage and subsequent mitochondrial β-oxidation, but when astrocytic mitochondria are compromised, this lipid buffering system fails, leading to pathological lipid accumulation and reactive astrocytes that secrete saturated lipids promoting neuronal injury. Experimentally, simply elevating neuronal membrane cholesterol is sufficient to reproduce early AD phenotypes, including enlarged endosomes, impaired vesicular trafficking, and increased Aβ productio^47^, while cholesterol metabolism has been identified as an independently druggable axis regulating both tau and Aβ in iPSC-derived AD neurons^48^. CP2 treatment re-enables this astrocyte-dependent lipid processing axis: the resulting normalization of cholesterol biosynthesis genes, increased cholesterol ester buffering, reduced sphingomyelin, and enhanced acylcarnitine shuttling observed in our data are consistent with a model in which CP2 achieves, from the mitochondrial side, a lipid metabolic rescue that both restores proper membrane composition across the neuron-glia unit and reduces the lipotoxic signaling that amplifies neurodegeneration. In this framework, CP2’s pleiotropic response is not a collection of independent pharmacological effects but rather an emergent property of a single upstream intervention (mild mtCI modulation) propagating through the interconnected network of mitochondrial bioenergetics, cholesterol homeostasis, membrane signaling, and astrocyte-neuron metabolic coupling.

The concurrent upregulation of *Smo, Notch*, and *EGFR* signaling components following CP2 treatment may reflect both a restoration of bioenergetic capacity and a normalization of membrane lipid composition required for receptor signaling competence. These pathways require substantial ATP for receptor trafficking, proteolytic processing, and downstream transcriptional activation, and their activity is sensitive to mitochondrial status. Mitochondrial dysfunction suppresses Notch signaling gene expression in neuronal cells^49^, while NAD⁺ restoration rescues Notch-dependent signaling in aged vasculature^50^. Similarly, EGFR signaling is tightly coupled to mitochondrial pyruvate metabolism through coordinated regulation of the glycolytic-TCA cycle. Beyond energetic constraints, however, membrane cholesterol itself is a potent modulator of G protein-coupled receptor (GPCR) function: the pool of "active" cholesterol that exceeds phospholipid binding capacity alters receptor behavior through both changes in bilayer biophysical properties and direct binding to transmembrane domains. Structural studies have identified specific cholesterol binding sites on GPCRs, including the β2-adrenergic receptor, and excess membrane cholesterol induces conformational changes that impair receptor activation and dimerization^51^.

Because norepinephrine and glutamate activate the astrocytic glycolysis underlying the ANLS through GPCR-mediated signaling^52^, cholesterol-driven impairment of these receptors provides a direct, non-energetic mechanism linking membrane sterol excess to disrupted metabolic coupling between astrocytes and neurons. In the AD brain, disrupted astrocytic glycolysis reduces lactate supply to neurons in a monocarboxylate transporter-dependent manner^14^, limiting the bioenergetic capacity to sustain these signaling networks. The recovery of *Smo, Notch*, and *EGFR* pathway gene expression alongside restored PDH flux, β-oxidation, and sphingolipid remodeling therefore, suggests that CP2 operates through a dual mechanism: the mitochondrial rescue expands the cellular energy budget while the consequent cholesterol normalization restores membrane biophysical properties required for GPCR-mediated neuron-glia communication, together re-engaging signaling programs that are silenced in the energy-depleted and cholesterol-overloaded AD brain.

To resolve the metabolic mechanisms underlying CP2’s pleiotropic effects, we developed *iMiceBrain*, the first genome-scale metabolic model of the mouse brain. Constructed as a consensus reconstruction from four pruning algorithms to maximize metabolic coverage, *iMiceBrain* demonstrated strong metabolic fidelity (80% essential task completion), high brain specificity (83–94% gene overlap with Allen Institute and HPA datasets), and significant translational potential with 91% of genes having identifiable human orthologs. Integration of experimental transcriptomics with *iMiceBrain* revealed that CP2 restores metabolic flux through the mevalonate pathway, particularly via Hmgcr-catalyzed reactions, predominantly through the Kandutsch–Russell pathway, the dominant route of brain cholesterol biosynthesis^53^. Transcriptomics confirmed reversal of APP/PS1-associated changes in cholesterol biosynthesis and clearance regulators, while lipidomics revealed elevated cholesterol esters and significantly reduced sphingomyelin following CP2 treatment, indicating a shift from pro-inflammatory sphingolipid signaling toward cholesterol storage and homeostasis. This result is supported by CP2-driven upregulation of *Sphk2* and enrichment for lipid storage and endosomal pathways, suggesting that CP2 redirects cholesterol toward buffered storage and ER-mitochondria transport to maintain homeostasis. Consistent with the previous experimental studies^54^, *iMiceBrain* identified CP2-driven reprogramming of mitochondrial bioenergetics. *In silico* flux simulations identified significant activation of the acylcarnitine-carnitine shuttle (ACRNTm) exclusively under CP2 treatment, facilitating fatty acid import for β-oxidation. Supporting this, lipidomics showed trends toward decreased triglycerides alongside increased fatty acids and acylcarnitines, consistent with enhanced lipid catabolism for energy production. This is a critical adaptation given that brain glucose hypometabolism, impaired TCA cycle activity, and neuroinflammation-driven microglial glucose consumption collectively limit neuronal energy supply in AD^38,55^. CP2 concurrently enhanced PDH activation (reduced pPDH and pPDH/PDH ratio), restoring pyruvate-to-acetyl-CoA flux and expanding the acetyl-CoA pool to sustain both oxidative phosphorylation and histone acetylation, including H3K27ac restoration. This dual activation of β-oxidation and PDH-dependent flux represents a compensatory, systems-level strategy resembling the adaptive fasting response, mobilizing lipid-derived and glucose-derived carbon to maintain ATP production and epigenetic regulation under chronic metabolic stress. Whether CP2 exerts parallel metabolic effects in peripheral organs such as the liver remains an important question for future investigation. While this study has focused on a short-term treatment in aged APP/PS1 mice, a previous study demonstrated that chronic CP2 treatment in female APP/PS1 mice starting at 9 to 24 months of age was safe and efficacious in blocking neurodegeneration and preserving cognitive function, suggesting the safety of such an approach^15^. Furthermore, recent studies using human organoids from LOAD patients carrying APOE4 allele demonstrated the efficacy of CP2 in reducing Aβ and pTau levels, suggesting this approach could work in patients with LOAD^24,25^.

Several limitations of this study should be acknowledged. First, our findings are derived exclusively from female APP/PS1 mice, a familial AD model carrying defined mutations. Whether CP2 elicits comparable effects in male mice or across diverse genetic backgrounds remains to be determined. Second, although *iMiceBrain* represents a significant advance as the first mouse brain-specific GEM, it assumes steady-state conditions and does not capture dynamic or compartment-specific metabolic regulation. Fourth, lipidomic and metabolomic trends supporting enhanced β-oxidation and pyruvate flux did not reach statistical significance, likely reflecting the small cohort size (*n* = 5). Future studies should evaluate the efficacy of mtCI modulators in both sexes across sporadic AD models, extend treatment duration to assess sustained disease modification, validate *iMiceBrain* predictions using isotope tracing or spatially resolved metabolomics, and characterize the peripheral metabolic footprint to fully delineate therapeutic potential.

In conclusion, this study demonstrates that CP2, a mtCI modulator, retains disease-modifying efficacy even when administered for a short time at advanced stages of AD-like pathology in aged female APP/PS1 mice. Through integrated transcriptomic, metabolomic, and lipidomic profiling, we show that CP2 elicits a coordinated adaptive response encompassing neuroprotection, restoration of proteostasis, normalization of circadian gene expression, and suppression of neuroinflammation, thereby reversing molecular signatures that define late-stage disease. Importantly, these findings establish a mechanistically validated therapeutic paradigm that extends beyond a single pathway. Ongoing studies with next-generation analogs, including C458^25^ and the clinically optimized lead C273^24^ demonstrate that the beneficial effects of weak mtCI modulation can be preserved and further enhanced through rational chemical optimization, with improved drug-like properties and translational readiness. Together, these results define a coherent drug discovery trajectory from proof-of-concept (CP2) to lead refinement (C458) and clinical candidate development (C273). Collectively, this work positions mtCI modulation and activation of adaptive stress responses as a robust and scalable disease-modifying strategy, with strong translational potential for AD and other age-associated neurodegenerative disorders^56^.

## Methods

### CP2 synthesis

Nanosyn, Inc. (http://www.nanosyn.com) synthesized CP2 as described previously^15^. Purification was conducted via high-performance liquid chromatography (HPLC), and NMR spectra were obtained for authentication and verification of drug purity. CP2 was made and stored as a free base, while aliquots of 20 µl volume were stored at –80°C, allowing its application in the study while avoiding freeze‒thaw cycles.

### Mice

Experiments with mice were performed following the approval of the Mayo Clinic Institutional Animal Care and Use Committee and in accordance with the National Institutes of Health’s *Guide for the Care and Use of Laboratory Animals*. IACUC Protocol A00001186-16-R24. This study was reported via the ARRIVE guidelines. The current research utilized mice carrying familial mutations in the *APP*(K670N/M671L) and *PS1*(M146L) genes (referred to as APP/PS1 mice) and their non-transgenic (NTG) littermates^17^. All the mice were females. PCR was used to verify the genotype, as described here^17^. Animal husbandry was performed under a 12–12 h light‒12 h dark cycle with a regular feeding schedule.

### CP2 treatment of NTG and APP/PS1 mice

NTG and APP/PS1 female mice (*n* = 5 mice/group) were given either CP2 (25 mg/kg/day in 0.1% PEG dissolved in drinking water *ad libitum*) or vehicle-containing water (0.1% PEG) starting at 19 months of age as described here^15^. The treatment lasted for six weeks. Mouse housing included five mice *per* cage, and weight and water consumption were monitored on a weekly basis to adjust the CP2 concentration. Random selection of mice was performed based on age and genotype, and the number of included mice was determined to achieve a 95% chance of detecting changes in 30–50% of the animals. The exclusion criteria for the study included significant (15%) weight loss, changes in grooming habits (hair loss), pronounced motor dysfunction (paralyses), or other visible signs of distress (unhealed wounds). None of the animals presented any of the above symptoms and were not excluded from the study. At the end of treatment, the mice were sacrificed, and brain tissue from the cortico-hippocampal region was collected for next-generation bulk RNA sequencing, Western blotting, and targeted metabolomics and lipidomics.

### Next-generation bulk RNA sequencing

Tissues from cortico-hippocampal brain regions were lysed via QIAzol (Qiagen cat. # 79306), and RNA extraction was performed via the miRNeasy (Qiagen cat. # 217004) according to the manufacturer’s instructions. RNA quantification and quality control were performed using a NanoDrop spectrophotometer and an Agilent 2100 Bioanalyzer, respectively, and all samples had an RNA integrity number (RIN) > 8.0. The RNA libraries were generated from 200 ng of RNA via the TruSeq RNA Library Prep Kit v2 (Illumina). RNA sequencing was performed at the Mayo Clinic Medical Genome Facility (MGF) Sequencing Core via the Illumina HiSeq 4000 platform with a paired-end 101-bp read length. For each sample, approximately 50 million single-fragment reads were obtained.

### RNA sequencing analysis and gene-set enrichment analysis (GSEA)

The RNA sequencing output was analyzed via MAP-RSeq v2.1.1, a comprehensive computational pipeline developed by the Mayo Clinic’s Division of Biomedical Statistics and Informatics, which applies publicly used bioinformatics packages. Through MAP-RSeq, we obtained gene counts, expressed single nucleotide variants (eSNVs), gene fusion candidates, and quality control plots. Read alignment and mapping were performed with TopHat2 against the mm10 reference genome. Gene and exon-level counts were obtained via feature Counts with Ensembl gene annotation files. The ‘-O’ option was applied to account for reads overlapping multiple genomic features, and the ‘-f’ option was used to quantify expression at the exon level.

Validation of data reliability and quality control monitoring were verified through plots created by RSeQC ^57^. Files with read counts were then processed for further analysis where the gene symbols were fetched via the bioconductor package biomaRt^58^ in R programming language v.4.2.2 by determining the attributes of “*external_gene_name*” and filters of “*ensembl_gene_id*”.

Moreover, protein-coding genes were captured by identifying the attribute “*transcript_biotype*”. Bulk RNA-transcriptomic data were first filtered to include protein-coding genes only. Second, genes with read counts <1 in at least three samples were excluded from further analysis. The Bioconductor package of Deseq2 ^59^ was used for performing differential gene expression (DEG) analysis, which was applied in the R programming language v.4.2.2. Genes were considered differentially expressed if they met the cutoff thresholds of an absolute log fold change (|log2 FC| ≥ 0.25) and a *p*-value < 0.01. DEG analysis was first performed considering the genotype as the main variant, allowing the profiling of APP/PS1 mice. Thereafter, treatment was included as a covariant, enabling the detection of the influence of treatment on the genotype. The set of differentially expressed genes was mapped against four libraries, namely, MGI (mammalian-phenotype)^60^, KEGG-human^61^, GO (biological-process) ^62,63^, and Reactome-2022^64^ using enrichr-KG^65^ to perform GSEA and identify the associated biological pathways.

### Targeted metabolomics

Mouse brain tissue was homogenized in 1× PBS using an Omni Bead Ruptor 24. Fifty-microliter aliquots of the homogenate were separated for metabolite quantitation. Targeted metabolite assays were performed on homogenates to quantify 11 tricarboxylic acid (TCA) cycle analytes and 8 short-chain fatty acids (SCFAs) by gas chromatography-mass spectrometry (GC-MS). In addition, 14 acylcarnitines (C0–C18:1), 12 nonesterified fatty acids (NEFAs), pyruvate, and acetyl coenzyme A (acetyl-CoA) were quantified via liquid chromatography–mass spectrometry (LC‒MS). Metabolite levels were normalized to the protein content of each sample and determined via the bicinchoninic acid (BCA) assay. Three-group comparisons were conducted via one-way ANOVA.

#### TCA cycle

The concentrations of the TCA analytes were measured via GC‒MS as previously described^66,67^ with a few modifications. Briefly, the homogenate was lysed via sonication and then extracted in 300 µl of chilled methanol and acetonitrile solution after the addition of 10 µl of internal solution containing U-13C-labeled analytes. After the supernatant was dried on a speed vac, the sample was derivatized with ethoxime and then with MtBSTFA + 1% tBDMCS (N-methyl-N-(t-butyldimethylsilyl)-trifluoroacetamide + 1% t-butyldimethylchlorosilane) before it was analyzed on an Agilent 5977B MSD GC/MS system under electron impact and single ion monitoring conditions. The concentrations of lactic acid (m/z 261.2), fumaric acid (m/z 287.1), succinic acid (m/z 289.1), ketoglutaric acid (m/z 360.2), malic acid (m/z 419.3), aspartic acid (m/z 418.2), 2-hydroxyglutaric acid (m/z 433.2), cis-aconitic acid (m/z 459.3), citric acid (m/z 591.4), isocitric acid (m/z 591.4), and glutamic acid (m/z 432.4) were measured against 7-point calibration curves that underwent the same derivatization procedure.

#### Acylcarnitine profile

Acylcarnitines (specifically C0–C18:1) were quantified via LC–MS as previously described^67,68^ with a few modifications. Briefly, each tissue homogenate was spiked with an isotopically labeled internal standard mixture containing acylcarnitines. The samples were extracted with cold methanol:dichloromethane (1:1, v/v) and centrifuged at 12,000 × g for 10 minutes. The resulting supernatant was transferred to a clean vial, evaporated to dryness, and reconstituted in running buffer. A calibration curve was prepared from a commercially available acylcarnitine mixture at various concentrations and spiked with the same internal standard used for the samples. All samples and calibration standards were analyzed on a triple quadrupole mass spectrometer coupled to an ultrahigh-pressure liquid chromatography (UHPLC) system. Data acquisition was performed in positive electrospray ionization mode via selected reaction monitoring (SRM). Each analyte was quantified by comparing its response to the corresponding calibration curve.

#### Non-esterified fatty acid (NEFA) profile

Non-esterified fatty acids (NEFAs) were quantified against calibration standards on a Thermo Quantis Plus triple quadrupole mass spectroeter coupled to a Thermo Vanquish Flex high-pressure liquid chromatography system (LC–MS), as previously described^69^. Briefly, each tissue homogenate was spiked with an internal standard prior to extraction with methyl tert-butyl ether (MTBE). The resulting extracts were dried under nitrogen and reconstituted in running buffer before LC–MS analysis. Data acquisition was performed in negative electrospray ionization mode.

#### Short-chain fatty acid (SCFA) profile

Short-chain fatty acids (SCFAs) were quantified by gas chromatography–mass spectrometry (GC–MS) as previously described^70,71^ with minor modifications. Briefly, homogenate was added to a tube containing 10 µL of an internal standard solution at pH 2 composed of isotopically labeled SCFA standards. A volume of three hundred microliters of dichloromethane (DCM) was used to extract SCFAs from the mixture. The extract was then derivatized with N-methyl-N-tert-butyldimethylsilyltrifluoroacetamide (MTBSTFA) prior to GC–MS analysis. Derivatized analytes were separated on a DB5MS column (30 m × 0.25 mm ID × 0.25 µm film thickness) and detected on an Agilent MSD5977A mass spectrometer. The concentrations of acetic acid (m/z 117.0), propionic acid (m/z 131.1), isobutyric acid (m/z 145.1), butyric acid (m/z 145.1), isovaleric acid (m/z 159.1), valeric acid (m/z 159.1), isocaproic acid (m/z 173.2), and hexanoic acid (m/z 173.2) were determined against 12-point calibration curves that underwent identical derivatization.

#### Acetyl-CoA quantitation

Acetyl-CoA was quantified by liquid chromatography–mass spectrometry (LC–MS) as previously described^72^. Briefly, an isotopically labeled acetyl-CoA internal standard solution was added to each homogenate. Proteins were precipitated by adding cold methanol, and the resulting supernatant was transferred to a new vial, dried and reconstituted in mobile phase A for analysis. A 12-point calibration curve was prepared using authentic acetyl-CoA standards spiked with the same internal standard solution as the samples. Both standards and samples were analyzed on a SCIEX 7500 triple quadrupole mass spectrometer coupled to a Nexera 40 liquid chromatography system. Data acquisition was performed in positive electrospray ionization mode via selected reaction monitoring (SRM) of transitions m/z 810 → 303 for acetyl-CoA and m/z 812 → 305 for [¹³C₂]-acetyl-CoA.

#### Pyruvate quantification

Pyruvate was quantified via the isotope dilution technique on an Agilent 6460 triple quadrupole mass spectrometer coupled with a 1290 Infinity II quaternary pump (LC‒MS/MS), as previously described^73^ with minor modifications. Briefly, a solution containing [^13^C_2_] pyruvate was added to the homogenate as an internal standard. Proteins were precipitated by adding cold methanol:acetonitrile (1:1, v/v) to the mixture. The supernatant was collected, dried, and reconstituted in running buffer prior to LC‒MS/MS analysis. Data acquisition was performed under negative electrospray ionization mode via multiple reaction monitoring (MRM) at m/z 87→45.1 and 89→43.1 for ^12^C-pyruvate and ^13^C_2_-pyruvate, respectively. Pyruvate concentrations were calculated from a 12-point calibration curve constructed from authentic standards.

Metabolomic analysis was conducted via the R package of MetaboAnalystR v4.0^74^, where the data were log transformed and scaled by mean centering. Differentially abundant metabolites were determined using the thresholds of a raw *p*-value < 0.05 and a fold change of 1.0 as cutoffs. A supervised multivariate analysis of oPLS-DA^75^ was conducted, followed by a permutation test with 10,000 iterations. The top 20 metabolites were identified as loadings for oPLS-DA.

### Untargeted metabolomics

#### Lipidomic profile

Lipids were extracted from homogenates with MTBE, dried and resuspended in the mobile phase prior to analysis on an LCMS system^76^. Data were acquired via an Orbitrap Exploris 120 mass spectrometer coupled with an UltiMate 3000 Binary Rapid Separation LC System (Thermo Fisher, USA). Lipids were separated chromatographically on an Accucore C30 HPLC column (2.6 µm, 2.1 × 250 mm; Thermo Fisher, USA). The analysis was performed in both positive and negative electrospray ionization modes, with full-scan coverage from m/z 200--1700 at 120K resolution.

Data-dependent acquisition targets the top 4 ions for fragmentation at 30K resolution. Subppm accuracy was maintained throughout the acquisition process via Thermo Fisher’s easy-IC internal calibrant.

The raw data files are imported into LipidSearch software for annotation and alignment MS peak areas are integrated, and lipids are identified by comparing MS/MS fragment ions with an internal database. After annotation, peak alignment combines the results from both positive and negative ionization modes. All the peaks were manually reviewed to prevent false identifications. The R package Lipidr^77^ was used for performing lipidomic analysis of the 907 lipid species detected. The data were processed for probabilistic quotient normalization (PQN) followed by unsupervised multivariate analysis via principal component analysis (PCA). Pairwise discriminative analysis of oPLS-DA was conducted for the treated groups. Differentially abundant lipid species were detected using the thresholds of *p*-value ≤ 0.05 and absolute log2-fold change (log2FC)| ≥ 1.0 as cutoffs. Differential lipids were processed for ontology enrichment analysis through mapping against the LION database.

### Reconstruction of genome-scale metabolic network of the mouse brain

A brain-specific genome-scale metabolic model (GEM) was reconstructed from the mouse model iMM1865 using four context-specific algorithms: mCADRE, rFASTCORMICS, CORDA, and ftINIT. We integrated bulk and total RNA-seq data from 93 mouse brain samples across 17 studies (NCBI-GEO) (**Supplementary Table 7**).

For mCADRE, iMM1865^32^ was validated against 31 essential metabolites prior to pruning, with gene expression ubiquity scores computed from limma-normalized data. For rFASTCORMICS and CORDA, TPM-normalized expression data were used. rFASTCORMICS fit dual Gaussian distributions to log₂(TPM) values per sample to derive zTPM scores, classifying genes as expressed (zTPM > 0; +1), not expressed (zTPM < −3.0; −1), or uncertain (0), which were then mapped to reaction penalties. CORDA classified reactions by confidence level using gene expression scores: NaN = 1, zero expression = 0, ≤25th percentile = +1, >25th–30th percentile = 2, and >30th percentile = 3. Metabolic task validation was performed throughout to ensure model functionality. For ftINIT, iMM1865 was first tested against essential metabolic tasks, then TPM-normalized data were integrated. Per-gene row-wise means were computed, and local 30th and 90th percentiles were averaged to define global lower (g_min) and upper (g_max) thresholds; genes were assigned Transcript Activity Scores of −1 (inactive), +1 (active), or 0 (intermediate). Pruning was performed in "1+1" mode, retaining reactions lacking GPR associations where necessary for functionality. mCADRE and rFASTCORMICS were implemented in MATLAB using the COBRA Toolbox v3.0^78^; ftINIT additionally required the RAVEN Toolbox v2.10.3^79^; CORDA was implemented in Python v3.9.6 via COBRApy v0.29.1^80^. LP and MILP problems were solved using CPLEX v12.10 and Gurobi v12.0.

A consensus model, *iMiceBrain*, was then constructed by removing blocked reactions from all drafts and retaining only reactions shared across all draft reconstructions covering shared metabolites. Glutamate transport ("GLUt6") and mitochondrial ATP synthase ("ATPS4mi") were designated as additional objective function reactions alongside the canonical "BIOMASS_reaction". ApolipoproteinE (apoE) was incorporated into iMiceBrain by introducing three apoE biosynthesis reactions ("r1112"), adopted from the Recon3D^29^ curated metabolic reconstruction, with gene-protein-reaction (GPR) associations mapped to *Apoe*; corresponding sink and demand reactions ("SK_HC00009_c" and "DM_HC00009_c", respectively) were added accordingly. An extracellular ATP demand reaction ("DM_atp_e") was also included. Gap filling^78^ was performed using iMM1865^32^ as the universal reference model to identify the minimal set of reactions required to sustain objective function flux. Functional annotations for genes, metabolites, and reactions were assigned via databases BiomaRt, PubChem, BiGG, and HMDB, supplemented by manual curation. Systems Biology Ontology (SBO) terms were annotated using the "annotateSBOTerms" function from the COBRA Toolbox v3.0^78^.

Model quality was evaluated using Memote v0.11.1 to assess dead-end metabolites, blocked reactions, verify stoichiometric, charge, and mass balance consistency. Model functionality was further benchmarked against a curated set of metabolic tasks adapted from prior studies^32,81,82^ in both MATLAB v2023a and Python v3.9.6 (**Supplementary Table 4**). For sanity checks, the upper bounds of 31 reactions were temporarily relaxed following reversibility verification against BiGG; reaction bounds were subsequently restored to match those in iMM1865^32^. Metabolite and reaction identifiers were also temporarily harmonized to satisfy the naming conventions required by the "performSanityChecksOnRecon" function. Finally, *in silico* single-gene knockout simulations were performed for a set of experimentally validated essential and non-essential genes (**Supplementary Table 5**) to quantify the impact of each knockout on optimal metabolic flux, using Flux Balance Analysis (FBA) and a linear implementation of Minimization of Metabolic Adjustment (lMOMA)^83^.

### Integration of transcriptome data with *iMiceBrain* and *in silico* metabolic flux simulation

To investigate the metabolic signatures associated with CP2 treatment, we integrated transcriptomics data with *i*MiceBrain using the iMAT (Integrative Metabolic Analysis Tool) algorithm in the COBRA toolbox v3.0^78^ in MATLAB 2019a, with the academic license of Gurobi Optimizer v12.0 to solve LP and MILP problems. Thereafter, Monte Carlo optimized Gibbs (OptGP) flux sampling was performed to examine the flux distributions across the subsystems in the context-specific reconstructions. The OptGPsampler^84^ was used with a thinning factor of 1000 to explore the entire solution space of feasible flux distributions that satisfy the metabolic model’s constraints, including the mass balance, reaction bounds, and objective function. The metabolic simulations were carried out in Python v3.9.7 while the COBRApy toolbox^80^ and Gurobi Optimizer v12.0 were used to solve the LP/MILP problems.

### Western blot analysis

Protein expression was assessed in cortico-hippocampal regions of vehicle- or CP2-treated NTG and APP/PS1 mouse brains (*n* = 5/ group) using Western blot analysis. Tissues were homogenized and lysed in 1× RIPA buffer (25 mM Tris-HCl, pH 7.6, 150 mM NaCl, 1% NP-40, 1% sodium deoxycholate, 0.1% SDS) supplemented with phosphatase inhibitor PhosSTOP (Roche, cat. #04906837001), protease inhibitor cocktail (cOmplete, Roche, cat. #11697498001), and serine protease inhibitor PMSF (Santa Cruz, cat. #sc-482875). Total protein lysates (20-25 µg) were resolved on 4-20% Mini-PROTEAN TGX™ Precast Protein Gels (Bio-Rad, cat. #4561096) and transferred to PVDF membranes (Bio-Rad, cat. #1620177). The following primary antibodies were used: pAMPK (T172) (40H9) (1:1000, Cell Signaling Technology, cat. #2535S, RRID: AB_331250); tAMPK (1:1000, Cell Signaling Technology, cat. #2532S, RRID: AB_330331); Acetyl-CoA Carboxylase (C83B10) (1:1000, Cell Signaling Technology, cat. #3676S); pACC (S79) (D17D11) (1:1000, Cell Signaling Technology, cat. #11818S); NRF2 (N2C2) (1:1000, GeneTex, cat. #GTX103322); Histone H3 (1:4000, Cell Signaling Technology, cat. #9715S); H3K27ac (D5E4) (1:1000, Cell Signaling Technology, cat. #8173S); β-Actin (1:50000, Sigma–Aldrich, cat. #A5316, RRID: AB_476743); GAPDH (14C10) (1:5000, Cell Signaling Technology, cat. #2118S); pULK1 (Ser555) (1:1000, Cell Signaling Technology, cat. #5869S, RRID: AB_10707365); and PGC1α (3G6) (1:1000, Cell Signaling Technology, cat. #2178S). Secondary antibodies included HRP-conjugated goat anti-rabbit IgG (H+L) (1:5000, Jackson ImmunoResearch, cat. #111-035-003) and HRP-conjugated goat anti-mouse IgG (H+L) (1:5000, Jackson ImmunoResearch, cat. #115-035-003). Signals were quantified using the ChemiDoc Imaging System (Bio-Rad), and densitometric analysis was performed using ImageJ software.

### Statistical analysis and data availability

Statistical analysis was conducted in R using the rstatix package within tidyplot^85^. All analyses were conducted via one-way ANOVA, followed by Tukey’s post hoc test. Statistical significance was determined following a *p*-value < 0.05. Differential gene expression analysis was performed following thresholds of |log2FC| > 0.25 and *p*-value < 0.01, whereas differential lipidomics was determined using thresholds of |log2FC| > 1.0 and *p*-value < 0.05. Kolmogorov‒Smirnov test was conducted in Python using scipy.stats module v1.16.2 for pairwise comparisons of reaction flux distributions in flux sampling, where statistical significance was set at *p*-value < 0.05.

## Data availability

Transcriptomics data is available in GEO database with the accession number GSE332871. Metabolomics and lipidomics data will be made available upon reasonable request to the corresponding author. The *iMiceBrain* model is available in BioModels with the following accession ID MODEL2606020001.

## Code availability

All the scripts and data used to develop the mouse-brain GEM (iMiceBrain) are available at (https://github.com/BaloniLab/iMiceBrain.git). Scripts used for omics analysis, figure generation, and integration of transcriptomics data with *iMiceBrain* are available at (https://github.com/BaloniLab/Tricyclic_Pyrone-CP2-.git) (last modified on May 26, 2026). The codes are also available in Zenodo (DOI: 10.5281/zenodo.20512211).

## Acknowledgments

This work was supported by National Institute of Health grants from the National Institute on Aging 4RF1AG055549-03 (to P.B. and E.T.); U19AG023122 (to N.R); UG3/UH3NS113776 (to E.T.); and the Alzheimer’s Association Research Fellowship grant 23AARF-1027342 (to T.K.O.N.). The funders had no role in study design, data collection and analysis, decision to publish, or preparation of the manuscript. The authors acknowledge the Mayo Clinic Medical Genome Facility Sequencing Core and Mayo Clinic Metabolomics Core for help with transcriptomic and metabolomics experiments. This manuscript was edited using Rubriq.

## Author contributions

**E.G.**: Data Curation, Methodology, Formal analysis, Investigation, Writing - Original Draft, Writing - Review & Editing. **T.K.O.N**: Experiments, Data Curation, Formal analysis, Investigation, Writing – Original Draft, Writing – Review & Editing. **T.K.:** Experiments, Formal analysis, Writing – Review & Editing. **H.G**.: Experiments, Formal analysis, Writing – Review & Editing. **N.R**.: Resources, Investigation, Writing - Review & Editing. **C.F.**: Investigation, Writing - Review & Editing. **P.B**: Conceptualization, Resources, Methodology, Formal analysis, Investigation, Writing - Original Draft, Writing - Review & Editing, Supervision. **E.T.**: Conceptualization, Oversight of experiments, Resources, Investigation, Writing - Review & Editing.

## Supplementary information

Supplementary Tables 1-7

Supplementary Figures 1-3

Supplementary File

